# Synergic Interaction between Permethrin and Bt toxins discovered using RNA-seq

**DOI:** 10.1101/684803

**Authors:** Takuma Sakamoto, Toshinori Kozaki, Norichika Ogata

**Affiliations:** Laboratory of Applied Entomology, Tokyo University of Agriculture and Technology, Tokyo, Japan; Laboratory of Applied Entomology and Zoology, Faculty of Agriculture, Ibaraki University, Ibaraki, Japan; Nihon BioData Corporation, 1-2-3 Sakado, Takatsu-ku, Kawasaki, Kanagawa, Japan

**Keywords:** Negatively correlated Cross-Resistance, Differential Expression, Digital Gene Expression, Permethrin, Bt toxins, Bombyx mori

## Abstract

Acting against the development of resistance to antibiotics and insecticides, involving negatively correlated cross-resistance (NCR) is an alternative to use- and-discard approach. It is termed NCR that toxic chemicals interact with each other and resistance of target organisms to one chemical is sometimes associated with increased susceptibility to a second chemical when; an allele confers resistance to one toxic chemical and hyper-susceptibility to another, NCR occurs. However, only 11 toxin pairs have been revealed to cause NCR in insects. Finding novel NCRs is needed for integrated pest management. We analyzed permethrin, an insecticide, induced transcriptomes of cultured fat bodies of the silkworm *Bombyx mori*, a lepidopteran model insect. Differentially expressed gene analyses suggested *Bacillus thuringiensis* (Bt) toxin was an NCR toxin of permethrin. NCR to permethrin and Bt toxins in *Thysanoplusia intermixta*, the agricultural pest moth, was examined; the children of permethrin survivor *T. intermixta* had increased susceptibility to Bt toxin. A novel NCR toxin pair, permethrin and Bt toxin, was discovered. The screening and developmental method for negatively correlated cross-resistance toxins established in this study was effective, in vitro screening using model organisms and in vivo verification using agricultural pests.

## I. Introduction

Insecticides are essential in food production and health protection. For instance, some 300 pesticides are in use to control about 900,000 different kinds of insects living within a global population of 7,400,000,000. Insecticides interact with each other and resistance of insects to one chemical is sometimes associated with increased susceptibility to a second chemical^1^. This interaction phenomenon is termed Negatively correlated cross-resistance (NCR). New information on NCR would be useful in integrated pest management. Up to now, several methods have been reported for screening and development of NCR toxins. However, only 17 toxin pairs have been revealed to cause NCR in insects (DDT and deltamethrina, DDT and phenylthioureab in *Drosophila melanogaster*, Bt toxins in *Plodia interpunctella*, Bt toxins Cry1Ac and Cry1F in *Helicoverpa zea*, Bt toxins in *Helicoverpa armigera*, Bt toxin and gossypol, Bt toxin and *Steinernema riobrave*, Bt toxin and *Heterorhabditis* bacteriophora in *Pectinophora gossypiella*, Bt and nucleopolyhedrovirus in *Plutella xylostella*, Pyrethroids and N-alkylamidesc, AaIT and pyrethroidsd in *Musca domestica*, AaIT and pyrethroidse in *Heliothis virescens*, Pyrethroids and diazinonf in *Haematobia irritans*, N-propylcarbamate and N-methylcarbamateg, Nephotettix cincticeps, Organo-phosphates and synthetic pyrethroids in *Tetranychus urticae*)^2^. Given that next generation sequencing technology has recently been developed, a plan to identify resistant genes by omics analyses and seek drugs which target resistant genes via a knowledge base has become feasible^3^. Thus, it would become possible to predict NCR toxins from omics analyses and then apply agents to induce resistance, and then use the next medicine for the children of survivors. In this study, permethrin was selected for omics analyses.

The moth, *Thysanoplusia intermixta* is a pest which feeds on *Asteraceae^4^* (e.g., burdock) and *Umbelliferae* (e.g., carrot). The genome of *T. intermixta* is unknown. Here, we selected a lepidopteran model insect, the silkworm *Bombyx mori*, for omics analyses. The silkworm genome was revealed in 2008^5^ and is well annotated. To avoid confusion with omics analyses after surgical removal of the guts and gut lumen, the primary cultured fat bodies of the silkworm in MGM-450 insect medium^6^ were used. Permethrin concentrations in the media followed a previous study where an appropriate permethrin concentration (0.25 mM) was obtained, resulting in biologically significant genes by referencing transcriptome diversity as the index of the extent of transcriptome changes^7^. To investigate the effects of permethrin on transcriptomes, we sequenced 6 transcriptomes from larval fat-body tissues exposed to permethrin. Freshly isolated tissues were cultured for 80 hours in MGM-450 insect medium, and then cultured for 10 hours in medium supplemented with 0.25 mM *cis*-permethrin or 0.25 mM *trans*-permethrin. The resistant gene was identified by differentially expressed gene analyses. Fortunately, the identified resistant gene was the target of the *Bacillus thuringiensis* (Bt) toxin^8^. The Bt toxin was predicted as a NCR toxin following the permethrin. For validation of the predicted NCR toxin, the probabilities of survival were compared for the natural population of *Thysanoplusia. intermixta* and the children of permethrin survivor *T. intermixta.* NCR to permethrin and Bt toxins, very common insecticides^9^, in *T. intermixta* is reported in this study. after fertilization.

## II. Materials and Methods

### Comparative Transcriptomics Establishment of Primary Culture

The p50 strain of the silkworm, *Bombyx mori*, was grown on fresh leaves of the mulberry, *Morus bombycis.* The larvae of the 5th instar were aseptically dissected 3 days after the 4th ecdysis and the fat body was isolated. More than 100 chunks of the tissue (approx. 2 mm^3^) were ex*cis*ed from the fat bodies of 24 larvae. Those tissue particles were incubated in cell culture dishes (ø = 35 mm; BD Biosciences, NJ, USA) with MGM-450 supplemented by 10% FBS (Biowest, Nuaillé, France) with no gas change. The tissue was cultured without antibiotics for 80 hours at 25°C. The infection of the microbes was checked by microscopic inspection. Infection-free tissues were used in the following induction assays.

### Chemicals

All the chemicals used in this study were of analytical grade. *cis*-permethrin (Wako Pure Chemical, Osaka, Japan) and *trans*-permethrin (Wako Pure Chemical, Osaka, Japan) were dissolved in acetone. This solution was diluted with three times its volume of ethanol just prior to mixing with the medium. The final concentrations of *cis*-permethrin and *trans*-permethrin were 0.25 mM. No antibiotics were used in the assays to maintain the primary culture.

### Induction Assay

The original medium was replaced with the *cis*-permethrin or *trans*-permethrin-containing medium in the induction assays. The primary culture tissues were incubated with 0.25mM *cis*-permethrin or *trans*-permethrin for 10 hours. The final concentrations of the inducers and the duration of incubation were determined by the previous report^7^. Inductions were terminated by soaking the tissues in 0.75 ml of TRIzol LS (Invitrogen, CA, USA), and the tissues were kept at −80°C until analyzed.

### RNA Isolation

Total RNA was extracted using TRIzol LS (Invitrogen) and the RNeasy Lipid Tissue Mini Kit (Qiagen, Hilden, Germany) following the manufacturers’ instructions as previously described^7^. Silkworm fat bodies (30 mg) soaked in 300 μL TRIzol LS were homogenized. We isolated total RNA from the homoginates as previously described^7^. The integrity of rRNA in each sample was checked using an Agilent 2100 Bioanalyzer (Agilent Technologies, Santa Clara, CA, USA).

### Library preparation and sequencing: RNA-seq

Sequencing was performed according to the TruSeq single-end RNA-sequencing protocols from Illumina for Solexa sequencing on a Genome Analyzer IIx with paired-end module (Illumina, San Diego, CA, USA). A total of 1 μg total RNA was used as the starting material for library construction, using the TruSeq RNA Sample Preparation Kit v2. This involved poly-A mRNA isolation, fragmentation, and cDNA synthesis before adapters were ligated to the products and amplified to produce a final cDNA library. Approximately 400 million clusters were generated by the TruSeq SR Cluster Kit v2 on the Illumina cBot Cluster Generation System, and 36–65 base pairs were sequenced using reagents from the TruSeq SBS Kit v5 (all kits from Illumina). Short-read data were deposited in the DNA Data Bank of Japan (DDBJ)’s Short Read Archive under project ID DRA006123.

### Data analysis and programs

Sequence read quality was controlled using FastQC program^10^. Short-read sequences were mapped to an annotated silkworm transcript sequence obtained from KAIKObase (http://sgp.dna.affrc.go.jp) using the Bowtie program. A maximum of two mapping errors were allowed for each alignment. Genome-wide transcript profiles were compared between samples. All the statistical analyses were performed using R software version 2.13.0. For identifying differentially expressed genes specifically from RNA-Seq data, we performed the differentally expression analysis based on a model using the negative binomial distribution called DESeq which is implemented as the DESeq package.

### Survival Analysis Worms for the survival analysis

*Thysanoplusia intermixta* larvae were originally collected near the Fuchu campus of Tokyo University of Agriculture and Technology at Fuchu, Tokyo and then successively raised on an artificial diet^11^ at 25 °C (L:D 16:8 h). Adult insects were fed a 10% sugar solution absorbed in cotton. A *T. intermixta* strain was selected from the parental native strain; pentamolting larvae feed: 1.5 cm^3^ artificial feeds soaked in racemic mixture 1.0× 10^-4^ *%* permethrin (Permite 5% Cream; Galderma Ltd., Lausanne, Switzerland) aqueous solution.

### The Survival analysis

Larvae were grown on an artificial diet at 25 °C (L:D 16:8 h). Eighty *T. intermixta* larvae were put into independent plastic cages (φ76×38 mm, approx. 90 mL), 40 larvae were the native strain and the other 40 were the permethrin-survivor strain. Twenty pentamolting larvae (10 larvae were native strain and 10 larvae were the permethrin-survivor strain) were exposed to Bt toxins (Toaro suiwa-zai CT; OAT Agrio Co., Ltd., Tokyo, Japan) from *Bacillus thuringiensis* serotype *kurstaki*, containing Cry IAa, Cry IAb and Cry I Ac; pentamolting larvae feed: 1.5 cm^3^ artificial feeds soaked in 1/50000, 1/60000, 1/70000 and1/80000 Bt toxins dissolved in water. A normal concentration of Toaro suiwa-zai CT is 1/1000 and it is 70ppm.

### Data Analysis

Statistical analysis was performed using R software version R 3.0.1^12^. We developed four models using the GLM function. The response variables were dead or alive of insects assumed to follow the binomial distribution. The explanatory variables were the concentrations of Bt toxins as continuous variables and the permethrin selection as categorical variables. First model is full model; the survival rates were explained by the interating two variables, the concentrations of Bt toxins and the permethrin selection. In second model, the survival rates were simply explained by the concentrations of Bt toxins and the permethrin selection. The survival rates were explained by only the concentrations of Bt toxins in third model and the survival rates were explained by only the permethrin selection in forth model. The Akaike information criteria (AIC) values for each model were 28.051, 26.058, 30.282 and 29.182, respectively. The second model was selected given that it had the lowest AIC value. Likely food ratio testing was performed between the first and second models. The LC50 were caliculated from the concentrations of Bt toxins and the insect survival rates using function pro.b in the library MASS.

## III. Results

Comparative Transcriptomics To identify resistant genes to *cis*-permethrin, a comparison was made between the transcriptomes of silkworm fat bodies cultured in in the medium without permethrin and the transcriptomes of silkworm fat bodies cultured in the medium containing 0.25 mM *cis*-permethrin. It was found that there were 30 statistically significant differentially expressed genes (FDR < 0.05) in the comparison data (Supplemental Figure 1, Table 1) using a statistical method based on a model using the negative binomial distribution, DESeq. In the case of the transcriptomic study using 0.25 mM *trans*-permethrin, it was found that there were 50 statistically significant differentially expressed genes (FDR < 0.05) (Supplemental Figure 2, Table 2). From the gene analysis, BGIBMGA013492, BGIBMGA010010, BGIBMGA009516, BGIBMGA009442, BGIBMGA008815, BGIBMGA007738, BGIBMGA006456, BGIBMGA006099, BGIBMGA005875, BGIBMGA004577, BGIBMGA004172 and BGIBMGA002119 were the biologically significant permethrin induced genes since these twelve genes were included in differentially expressed genes in both comparison studies. The differentially expressed gene (BGIBMGA007738) codes ABC transporter C family protein. ABC transporters play a main role in the mode of action of Bt Cry toxic proteins^13–15^. We hypothesized Bt toxin is a NCR toxin following the permethrin; Permethrin exposed insect increases the expression of the ABC transporter and the transporter is a target of Bt toxin.

**Table 1.**
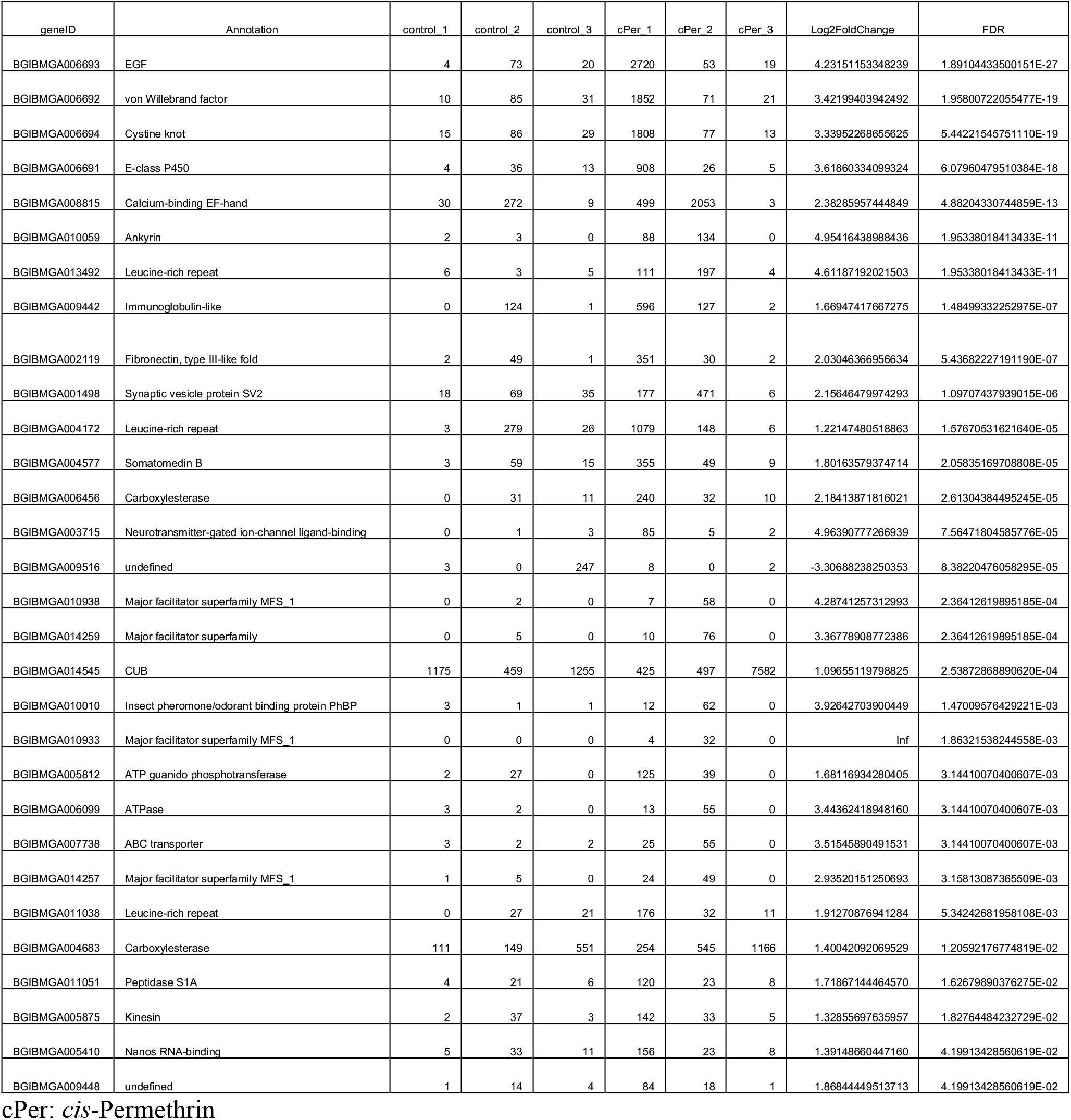
Differentially expressed *Bombyx mori* genes exposed to *cis*-permethrin

**Table 2.**
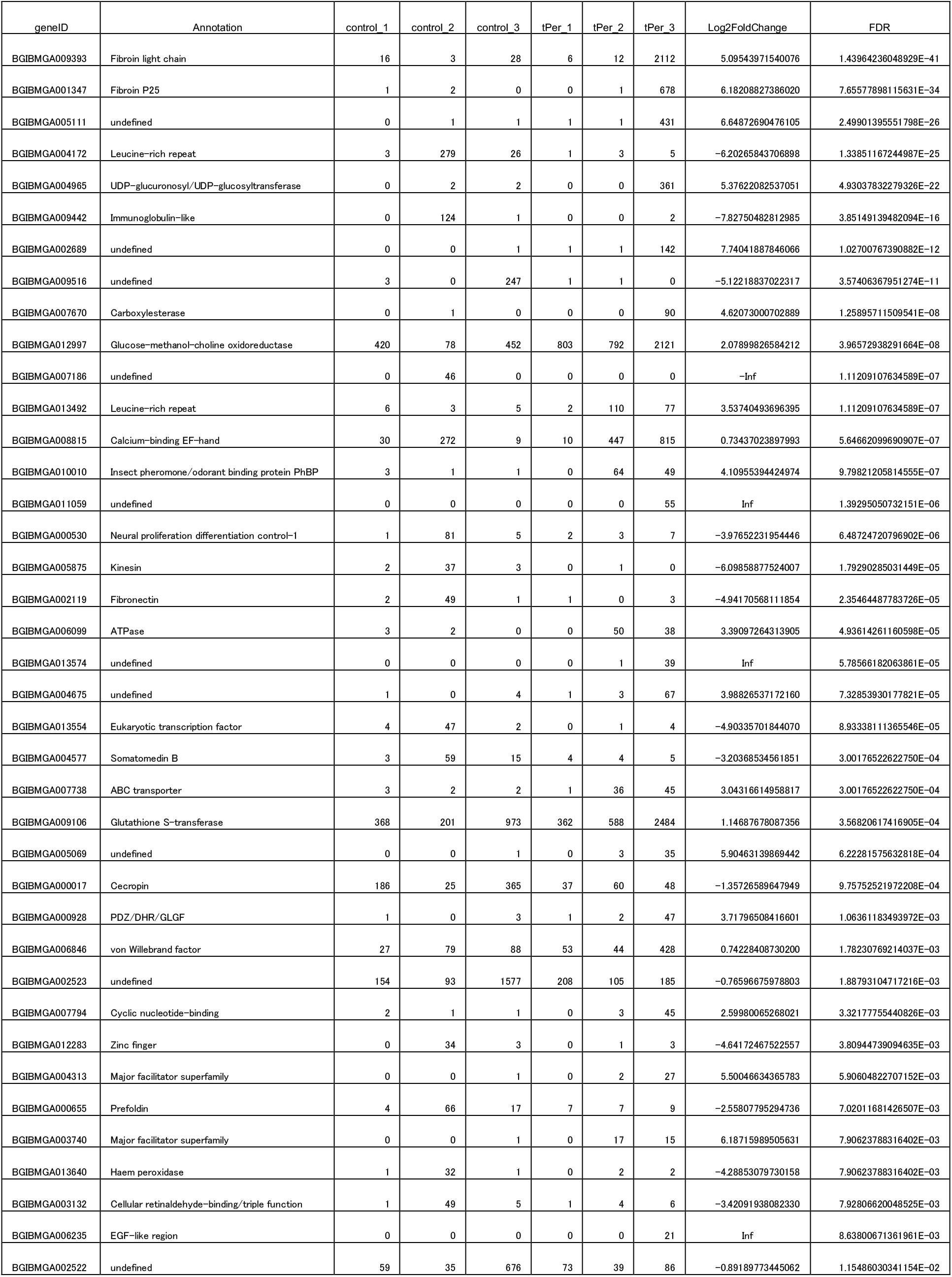
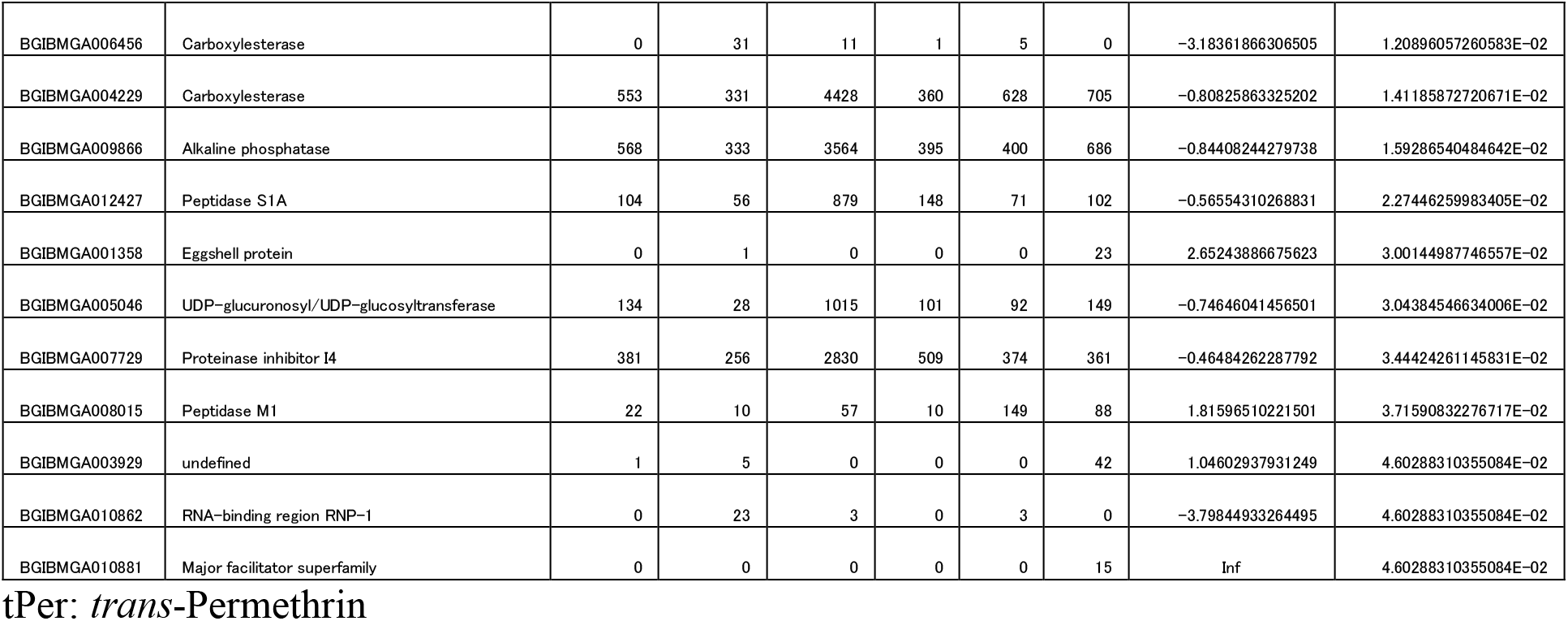
Differentially expressed *Bombyx mori* genes exposed to *trans*-permethrin

### Survival Analysis

The probabilities of survival for the native strain of *T. intermixta* and the children of permethrin survivors *T. intermixta* were compared. A *T. intermixta* strain was selected from the parental native strain using artificial feeds containing the permethrin. Each of the 10 insects of both strains was fed the artificial feeds containing the Bt toxins at 1/50000, 1/60000, 1/70000 and 1/80000 dilutions. The probabilities of survival of each insect were statistically analyzed using general linear models (GLMs). The permethrin selection was statistically significant (p = 0.0126), in a likelihood ratio testing between the selection containing model and the selection ignoring model. Lethal dose 50 (LC_50_) were 1.83 ppm in the native strain and 1.42 ppm in the permethrin-survivor strain.

## IV. Discussions

Negatively correlated cross-resistance to permethrin and Bt in *Thysanoplusia intermixta* was predicted using comparative transcriptomics of primary cultured fat bodies of a lepidopteran model insect, the silkworm *Bombyx mori.* Next generation sequencing technologies that reveal many genomes of non-model organisms and the annotating of genomes still represents a challenge. Recently, effective genome annotating tools were developed using genomes including the annotations of other organisms (https://github.com/Hikoyu/FATE^16^). Well-annotated genomes of model organisms are strongly desired. In this context, there are many harmful insects and NCR studies are needed for all harmful insects. However, the costs of insect species-specific screening and developing NCR strategies are prohibitive. Hence, screening and development programs for NCR toxins using model organisms should be initiated.

The transcriptomes from primary cultured tissues were sequenced in this study. Tissue cultures were devised as a means for studying the behavior of animal cells in an environment that is free from systemic variations that might arise *in vivo* both during normal homeostasis and under the stress of an experiment. In transcriptome analyses, p-values from a comparison of one replicate with another one must be below 0.05 in comparative transcriptomics with Type-I error control. Between replicates, no genes are truly differentially expressed, and the distribution of p-values is expected to be uniform in the interval [0,1]^17^ Whole larval transcriptome analyses would be compromised by the gut lumen. Surgical gut removal would result in changes in the volume ratio between tissues in a body. Differences from these variable factors would be obtained causing statistically significant differences between samples. RNA samples were readily obtained from cultured tissues in the present study thus avoiding such variable factors; in this way, biologically significant differentially expressed genes were realized. Drug concentration is also an important factor in comparative transcriptomics as has been reported^7,18,19^. Such methods, predicting NCR toxins using comparative transcriptomics, would be useful not only in pest control but also in research fields affected by biological drug resistance, e.g., antibiotics and cancer research.

## V. Conclusion

NCR is a strategy for acting against the development of resistance to insecticides. However, there is only 11 NCR toxin pairs have been revealed in insects. Here, we reported the novel NCR toxin pair, permethrin-Bt toxin. *In vitro* transcriptome analyses using cultured fat bodies of silkworm *bombyx mori*, a lepidopteron model insect, were performed to predict the NCR toxin pair. The predicted toxin pair was verified using *in vivo* crossresistance analyses using larvae of *Thysanoplusia intermixta*, the agricultural pest moth. In *in vitro* transcriptome analyses, we cared drug concentration to perform the perfect comparison. In genomic scale analyses, statistical significant differences must appear in any comparison. Comparison designs concentrate biologically significant differences in statistical significant differences.

**Fig 1.**
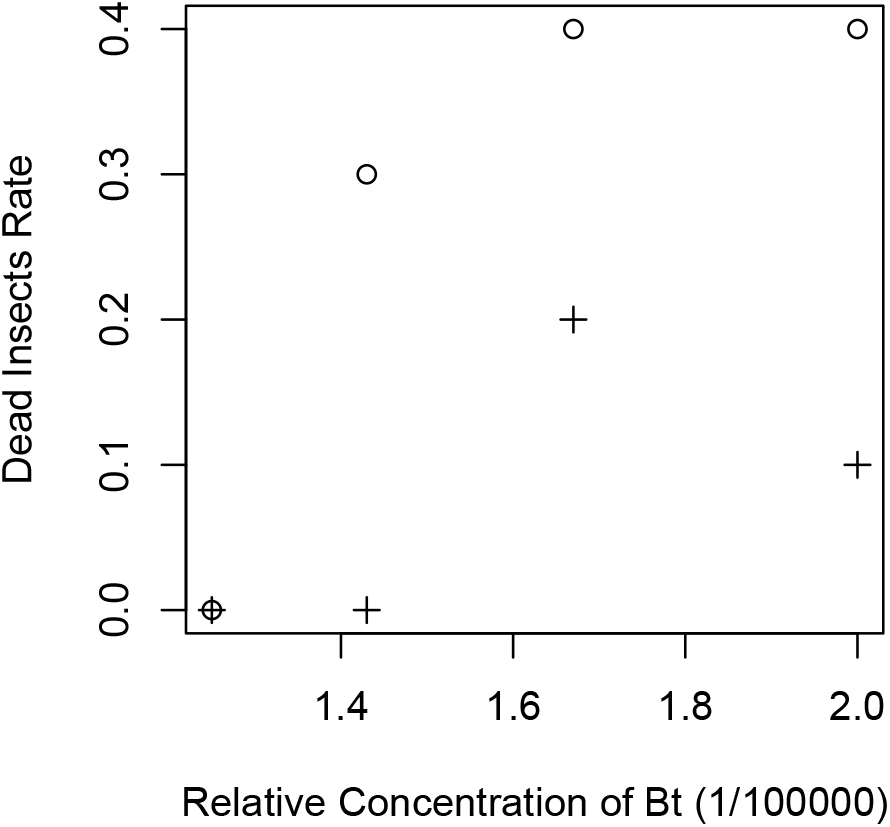
Probabilities of Survival of the Permethrin Survivor Strain and the Native Strain of *Thysanoplusia intermixta*. Scatter plot for rate of formation of dead insects versus Bt toxin concentration. The insects of the permethrin survivor strain are represented by O and the insects of the native strain are represented by +. The rates of formation of dead insects were estimated using 10 insects for each category. A total of 80 insects were used.

---

**Supplemental Figure 1.**
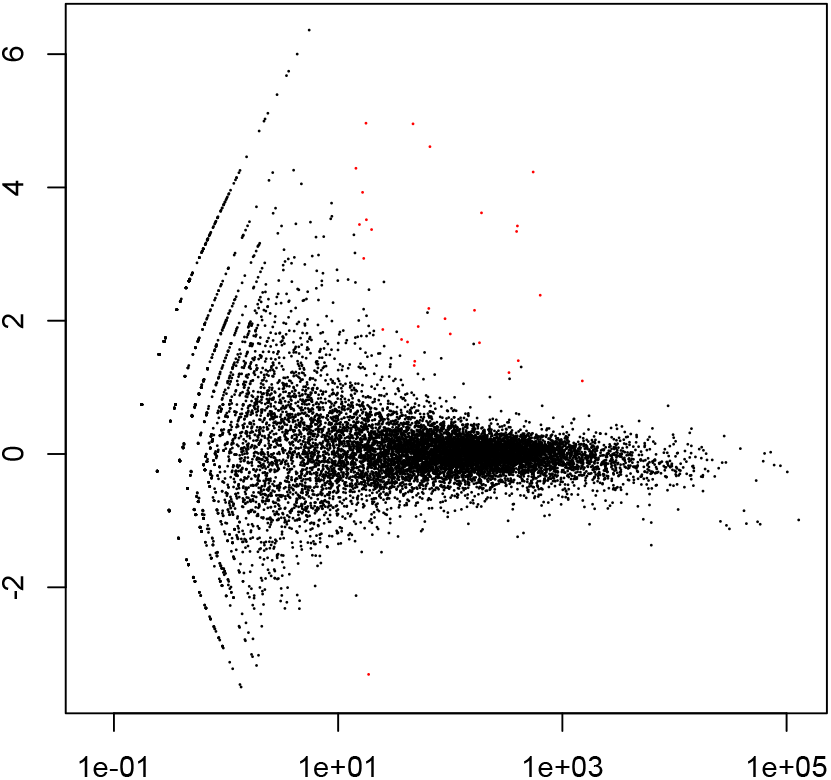
Differentially expressed gene analysis of *cis*-permethrin exposed cultured fat body of *Bombyx mori.* Scatter plot of log2-fold changes versus mean. Genes marked in red were detected as differentially expressed at the 5% false-discovery rate (FDR) using DESeq. A=log2((cPer/Ncp*control/Nco)^1/2^), M=log2(cPer/Ncp)-log2(control/Nco). “cPer” means expression value of genes in each cPer sequence data. “Ncp” means sum of expression value of genes which measured in each cPer sequence data, “control” means expression value of genes in each control sequence data. “Nco” means sum of expression value of genes which measured in each control sequence data.

**Supplemental Figure 2.**
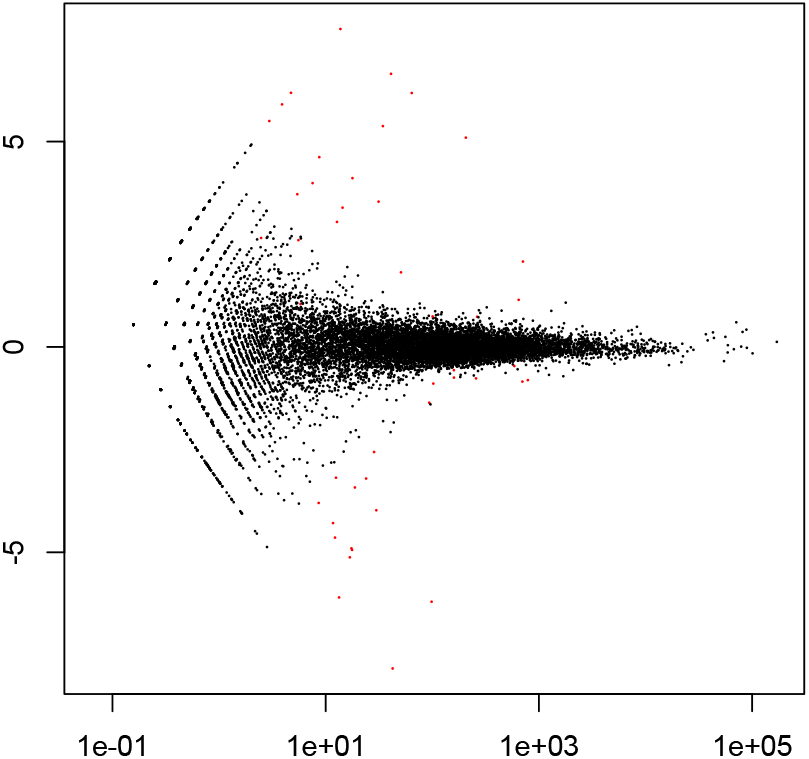
Differentially expressed gene analysis of *trans*-permethrin exposed cultured fat body of *Bombyx mori.* Scatter plot of log2-fold changes versus mean. Genes marked in red were detected as differentially expressed at the 5% false-discovery rate (FDR) using DESeq. A=log2((tPer/Ncp*control/Nco)^1/2^), M=log2(tPer/Ntp)-log2(control/Nco)

